# Group size and modularity interact to shape the spread of infection and information through animal societies

**DOI:** 10.1101/2021.05.01.442253

**Authors:** Julian C Evans, David J Hodgson, Neeltje J Boogert, Matthew J Silk

## Abstract

Social interactions between animals can provide many benefits, including the ability to gain useful environmental information through social learning. However, these social contacts can also facilitate the transmission of infectious diseases through a population. Animals engaging in social interactions must therefore face a trade-off between the potential informational benefits and the risk of acquiring disease. In order to understand how this trade-off can influence animal sociality, it is necessary to quantify the effects of different social structures on individuals’ likelihood of acquiring information versus infection Theoretical models have suggested that modular social networks, associated with the formation of groups or sub-groups, can slow spread of infection by trapping it within particular groups. However these social structures will not necessarily impact the spread of information in the same way if its transmission is considered as a “complex contagion”, e.g. through individuals copying the majority (conformist learning). Here we use simulation models to demonstrate that modular networks can promote the spread of information relative to the spread of infection, but only when the network is fragmented and group sizes are small. We show that the difference in transmission between information and disease is maximised for more well-connected social networks when the likelihood of transmission is intermediate. Our results have important implications for understanding the selective pressures operating on the social structure of animal societies, revealing that highly fragmented networks such as those formed in fission-fusion social groups and multilevel societies can be effective in modulating the infection-information trade-off for individuals within them.

**Significance statement:** Risk of infection is commonly regarded as one of the costs of animal social behaviours, while the potential for acquiring useful information is seen as a benefit. Balancing this risk of infection with the potential to gain useful information is one of the key trade-offs facing animals that engage in social interactions. In order to better understand this trade-off, it is necessary to quantify how different social structures can promote access to useful information while minimising risk of infection. We used simulations of disease and information spread to examine how group sizes and social network fragmentation influences both these transmission processes. Our models find that more subdivided networks slow the spread of disease far more than infection, but only group sizes are small. Our results demonstrate that showing that fragmented social structures can be more effective in balancing the infection-information trade-off for individuals within them.

## Introduction

Most animals engage in some form of social interaction (Krause et al. 2002; Frank 2007; Clutton-Brock 2016). These interactions frequently provide opportunity for gaining social information (Danchin et al. 2004; Laland 2004; McGregor 2005; Allen et al. 2013; Aplin et al. 2015), whether it be the location of new food sources (Aplin et al. 2012; Kendal et al. 2015; Berdahl et al. 2018; Nöbel et al. 2018) or predation risk (Beauchamp et al. 2012; Crane and Ferrari 2013; Frechette et al. 2014). However, interacting with others can also be costly, with one of the primary costs being the risk of infection with pathogens or parasites (Daszak et al. 2000; Stattner and Vidot 2011). Balancing risk of infection with the potential to gain useful information is therefore one of the key trade-offs facing animals that engage in social interactions (Evans et al. 2020; Romano et al. 2020). In order to better understand this trade-off, it is necessary to quantify how different social structures can promote access to useful information while minimising risk of infection (Evans et al. 2020; Firth 2020; Romano et al. 2020). Individuals occupying particular positions in these social structures can have higher fitness (Cameron et al. 2009; Wey et al. 2013) and certain social structures can also help maximise the benefits of sociality for their members, promoting the persistence of social units (Ilany and Akcay 2016; Kramer and Meunier 2019). Therefore social behaviours that influence emergent social structures will be target for selection, and finding the aspects of sociality that promote the spread of disease or information represents an important advance that contributes to our understanding of social evolution.

Animal social systems vary widely among species (Krause et al. 2002). Social networks offer a useful toolkit to capture and quantify this diversity (Krause et al. 2007; Wey et al. 2008; Pinter-Wollman et al. 2014), and can help distinguish species that, for example, live in multi-level societies (VanderWaal et al. 2014; Cantor et al. 2015; Papageorgiou et al. 2019), stable social groups (Weinrich 1991; Wey et al. 2013; Shizuka et al. 2014), fission-fusion groups (Kerth and Konig 1999; Couzin and Laidre 2009; Silk et al. 2014), or whose interactions and associations are more constrained by other factors such as shared space or resource use (Davis et al. 2015; Spiegel et al. 2016; Evans and Morand-Ferron 2019). Social network structure plays a critical role in governing both pathogen and information transmission (Aplin et al. 2012; Godfrey 2013; Webster et al. 2013; Silk et al. 2017; Evans et al. 2020). For example, variability in the connectedness of individuals can drive a superspreader effect in infectious diseases, causing more explosive outbreaks (Lloyd-Smith et al. 2005). Global properties of the network such as the number (or density) of connections and how these are distributed are also important. The efficiency of information transfer is maximised when social networks show intermediate levels of subdivision into different modules or communities (Romano et al. 2018). Many animal social networks possess such modular structures (Wey et al. 2008), especially species that form relatively stable social groups (Drewe et al. 2009; Weber et al. 2013). Consequently, the role of both community structure and how connected these communities are – their modularity – are likely to be important in governing any differences between information and disease transmission in animal societies.

There are often important differences in how infection and information spread through a network. Infection is generally considered a simple contagion (Moore and Newman 2000); the likelihood of becoming infected will depend predominantly on the number of relevant contacts with infected individuals, and the duration of these contacts (Godfrey et al. 2009). In contrast, information can often be considered a complex contagion (Macy 1991; Centola 2010; Firth 2020); individuals decide on whether to act on the information acquired and so may use different social learning strategies that change how information spreads (Laland 2004; Kendal et al. 2018). For example, they might accept information only from certain individuals (e.g. Kavaliers et al. 2005; van de Waal et al. 2010), or be much more likely to use it if a set proportion of their contacts behave in that way(Danchin et al. 2018). The latter can result in conformist social learning, which has been demonstrated in taxonomically diverse animal societies (van de Waal et al. 2013; Aplin et al. 2015; Danchin et al. 2018).

Because of the differences in how they spread, modularity shapes the transmission of infection and information in different ways (Nematzadeh et al. 2014; Sah et al. 2017a; Evans et al. 2020). While modular networks can promote the spread of infection within particular groups, they tend to trap infection within these groups, slowing down how quickly disease can spread through the population as a whole (Sah et al. 2017a). The impact of modular structure on disease outbreaks is greatest when the modularity of these subdivisions is high (i.e. different groups have relatively few contacts between them; Salathé and Jones 2010b; Sah et al. 2017a), especially when transmissibility is low so that infection is unlikely through occasional or rare contacts (Griffin and Nunn 2012; Sah et al. 2017a; Rozins et al. 2018). In contrast, when information transmission is a complex contagion, modular networks can promote the global spread of information, especially at intermediate modularities (i.e. when contacts between groups or modules are infrequent but not rare) due to strong social reinforcement within groups (Nematzadeh et al. 2014). As a result, modular networks may provide one route to promote the spread of information through a population disproportionately to the spread of infection (Evans et al. 2020). However, there are many other aspects of the structure of modular networks that might also shape this pattern and have been less well studied.

Social group (or sub-group) size, which is highly variable across and within animal societies, represents one such trait that could also impact how network structure shapes transmission dynamics (Côté and Poulinb 1995; Nunn et al. 2015). Sah et al. (2017a) showed that modular networks caused the greatest reduction in epidemic spread when network fragmentation was high, i.e. networks were composed of many smaller groups. This suggests that the formation of smaller social groups is more effective in reducing infectious disease transmission. However, different levels of network fragmentation may also change the optimum modularity for information transmission, which depends on high clustering of connections within groups to promote the spread of information in the first place (Nematzadeh et al. 2014). For example, small modules or sub-groups that contain many friends of friends may help promote the spread of information while limiting infectious disease transmission if between group contacts are rare. This could be especially important as the impact of modularity on infection transmission is non-linear, with no substantial effect on disease spread when modularity is below a certain value (Sah et al. 2017a). Thus, if the optimal level separation between groups for information spread is too far below the level at which a modular network structure begins to protect effectively against the spread of infectious disease then the benefits are lost.

We use simulation models to examine how social group size (or network fragmentation) interacts with modularity (the strength of the subdivision of a network into groups) to determine differences in the speed of social information spread and infectious disease transmission. We treat the spread of information using a conformist model in which the relationship between the likelihood of acquiring/using information and proportion of informed contacts is sigmoidal while infection risk is fixed for each contact with an infected individual. We compare the rate of spread of infection and information through social networks that vary in both their modularity and the size of groups making up the modular structure of the network, for a range of different levels of social connectedness and transmissibility values (how likely a disease or a piece of information is to be transmitted). We expect that modular networks would promote the transmission of information relative to the spread of infection, especially when transmissibility is low. However, we anticipate that the extent of this benefit may also be shaped by the number and size of social groups.

## Methods

### Simulation objectives

We generated simple networks with different edge density (the social connectedness of individuals), group size and modularity and used them to examine how these properties affect the transmission rates of information and disease, at varying levels of transmissibility.

### Simulation summary

Each simulation consisted of two main steps. First we generated random networks which were used to select parameters controlling disease and information spread. We then altered these random networks to create modular networks, in which we compared relative transmission speeds.

We generated a random network of 200 nodes (i.e. individuals) with a particular edge density (proportion of potential pairs of individuals in the network who were connected; Table 1). We then simulated the spread of a disease through this network, starting from a randomly selected individual, using a susceptible-infected (SI) model with the transmissibility of that disease in that network derived from an adjusted, per time interval R_0_ (Table 1). Disease simulation was then carried out in the random network 50 times, each time starting with a different randomly selected individual. For each simulation we recorded the number of timesteps it took a disease to infect 75% of the individuals in that network., We used the distribution of timesteps taken to reach this 75% threshold to determine the value of a scaling parameter γ (see below) so that information spread at a similar speed to disease in the random network. We simulated information spread using this parameter in the random network 50 times, starting with a random node, and recorded the average time taken for 75% of individuals to be informed.

**Table 1:** Summary of the four parameters used in the simulations

| Parameter | Description | N. values | Values |
| --- | --- | --- | --- |
| Network density | **Proportion of potential connections existing in the network.**<br>Higher densities result in more edges in the initial random network and in derived modular networks. | 9 | 0.02, 0.03, 0.04, 0.05, 0.06, 0.07, 0.08, 0.09, 0.1 |
| $R_0$ | **Basic reproduction number, infectiousness of disease.**<br>Used to derive values of $\beta$ (transmission probability in a particular random network) and $\gamma$ (scaling parameter to ensure information spreads at the same rate as disease in that random network). | 5 | 1.1, 1.25, 1.5, 2, 3 |
| Group size | **Social group size.**<br>Controls the size of social groups in the modular networks derived from a random network. | 6 | 5, 8, 10, 20, 25, 40 |
| Network modularity | **Strength of division of a network into modules.**<br>Larger values usually mean less intergroup edges in the modular networks derived from a random network. | 3 | 0.4, 0.6, 0.8 |

We then rewired the edges of the random network to produce new modular networks with the same edge density but a particular network modularity, with individuals subdivided into groups of a particular size. This led to a total of 18 modular networks being derived from each random network, one for every combination of modularity and group size (table 1). We simulated the spread of disease and information through these new modular networks and recorded the average time each process took to infect or inform 75% of nodes in the rewired networks. We then compared the difference in mean time taken for information and infection to reach the 75% threshold in that particular modular network. We also compared the transmission rates through a modular network to those through the random network from which it was derived, to establish the relative effect of modularity and group sizes on the transmission speeds of the two spreading processes.

The values of network density, transmissibility, group size and network modularity tested led to a total of 810 parameter combinations (table 1), with each combination repeated 20 times. We conducted all simulations in R 4.0.2 (R Development Core Team 2020). R code is provided in the Supplementary Materials. See supplementary video 1 for a visualisation of these simulations.

## Detailed methods

### Spread-simulation overview

We here summarise the methods used to simulate information and disease transmission throughout these simulations. We modelled both disease and information transmission as simple susceptible-infected (SI) models. Nodes were therefore either susceptible (uninfected/uninformed) or infected (informed), with no possibility for recovery from this state.

### Disease-spread simulation

When simulating a disease spread, an individual’s likelihood of infection will depend on the infectiousness of the disease and how many of their contacts are infected. We begin by infecting a random node (i.e. individual) in the network. In each subsequent timestep, whether a susceptible node will become infected is determined by a binomial trial, with the probability of infection for a node *v* being:

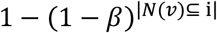

Where | *N*(*v*) ⊆ I | is the number of adjacent nodes to vertex *v* (*N*(*v*)) that are infected. *β* is a per timestep transmission probability per connection for a particular random network (see below). The likelihood of a node becoming infected therefore increases as the number of infected connections increases (See supplementary figure 1a).

### Information-spread simulation

We use a conformist learning rule to simulate the spread of information. Under this rule, an individual’s likelihood of accepting the information will depend on a scaling parameter and the proportion of contacts who have already accepted the information, with the probability of accepting increasing once 50% of contacts have accepted the information. The likelihood of accepting the information based on the proportion of contacts who have accepted the information is therefore a sigmoidal function, described in the equation below (see Fig. S1). A random node starts with the information, and in each subsequent timestep whether a susceptible node *v* will accept the information and become informed is based on a binomial trial, with the probability of accepting the information being:

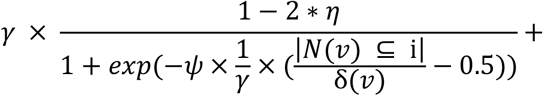

Where *γ* is a scaling parameter (see below for parameter selection process), | *N*(*v*) ⊆ I | is the number of adjacent nodes that are informed and δ (*v*) is the node’s degree. is the base probability of accepting information and *ψ* is the steepness of the sigmoidal function (See supplementary figure 1b). Based on our exploration of parameter values to match the rate of spread of information to the rate of disease spread in the random networks (see below), we fixed *η* at 0.001 and *ψ* at 10 in all simulations.

### Random network generation and parameter selection

We generated random networks consisting of 200 nodes using the igraph’s implementation of the Erdos-Renyi model (Erdős and Rényi; Csardi and Nepusz 2006). Probability of edge formation between nodes was determined by the value of the density parameter under investigation. Networks were binary (i.e. individuals were connected or not) and undirected (i.e. either both or neither individual were connected to each other). We then modelled the spread of a disease of each value of transmissibility, R_0_ (see table 1), through this random network. This allowed us to establish a baseline disease spread speed in a random network of this density, which could later be compared to the spread speed in modular networks. We also used this spread of disease to choose parameters allowing information to spread at the same rate.

### Disease spread parameter selection

To simulate the spread of disease through a network, we converted a chosen per-time period R_0_ to a transmission probability for the random network under investigation. First, we calculated *r*, the transmission probability for the period of infection *t* in that network, where *t* is the time period over which an R_0_ is calculated, which was fixed at 100. For example, for an R_0_ of 3, an individual should on average infect 3 other individuals in this network over 100 timesteps.

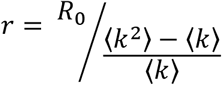

Here ⟨ *k* ⟩ is the average degree of all nodes in the network and ⟨ *k* ^2^ ⟩ is the average squared degree of all nodes in the network (Newman 2008). We then used the resulting value of *r* to calculate *β*, the per timestep transmission probability per connection for this R_0_ in this network.

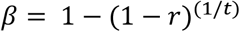

We used this value of *β* when simulating disease spread through the random network under investigation and for all modular networks derived from that initial random network, for this value of R_0_ (Newman 2008).

### Learning rule parameter selection

Having established the value of that corresponded to a value of R_0_ in the current random network, we then found the equivalent parameter value that lead to a similar transmissibility for information. To do this, we first simulated the spread of disease through the random network 50 times, recording the number of timesteps it took to infect 75% of the nodes. We used this distribution as a baseline with which to compare the spread of information. We searched for a value of a scaling parameter that governed the speed of information transmission, *γ*, which would produce a similar distribution of information spread (time to inform 75% of nodes) as our infection baseline. When generating a distribution, information spread through the random network was simulated 50 times with a potential value of *γ*, using the learning rule described below. We then compared the distribution of timesteps from these 50 simulations of information spread to that of the 50 simulations of the spread of disease by calculating the Bhattacharya distance (Bhattacharyya 1946) between the two distributions, where smaller values indicate a greater overlap between them. We optimised *γ* so as to minimise the Bhattacharya distance between the distribution of disease and information spread times using R’s “optimise” function (Nelder and Mead 1965). Parameter optimisation was run 5 times so as to avoid being trapped in local optima based on potential combinations of starting nodes and potential *γ*. From these five parameter searches, we selected the value of *γ* that produced the smallest Bhattacharya distance, (i.e. resulted in information spreading at a similar speed to disease) and recorded the mean time taken for each to spread to 75% of individuals in the network.

### Modular network generation and comparisons

We then altered the network to test how modularity and group sizes affected these spreading processes. For each random network we generated 18 modular networks, varying in group size and modularity. The random network was rewired to generate community structure with the desired group size and relative modularity (Q_rel_). Q_rel_ adjusts the modularity calculated by the maximum obtainable modularity with the same number of nodes and edges in the network (Sah et al. 2017b). The algorithm first assigns every node to a group, with the number of groups depending on the desired group size. It then rewires the network one edge at a time until the relative network modularity reaches its target. See Figure 1 for an example of the modular networks generated for different densities and group sizes.

**Figure 1:**
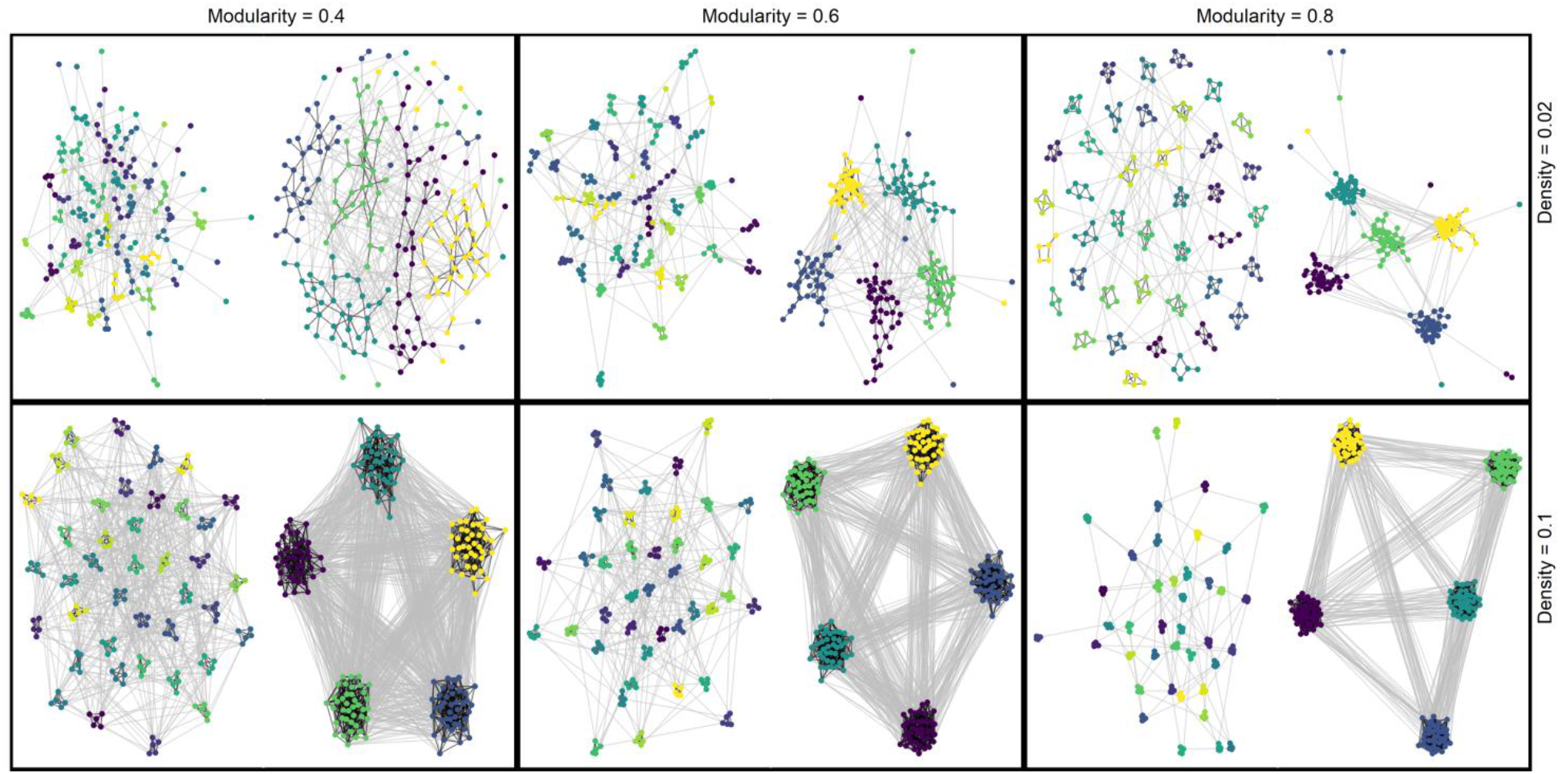
Examples of the networks at different levels of network density (row), rewired to different values of network modularity (column) and two different group sizes (5 and 40) for the same population size (200).

We then simulated information and disease spread through each of these new modular networks 50 times, using the parameters β and γ derived from the random networks. We recorded the mean time it took for 75% of nodes to become infected and the mean time it took for 75% of nodes to become informed in each modular network, with a cut off if one or both of the spreading processes failed to reach this threshold after 3500 timesteps. See supplementary video 1 for an example of simulations in both random and modular networks of varying densities.

## Results

### Summary of main results: the importance of modularity and group size

Speed of transmission of both disease and information was reduced by modular network structure, but mainly when group sizes were smaller and so networks more fragmented (Fig. 2). Higher levels of transmissibility always led to faster spread of disease and information, in all types of modular networks. However, more modular and fragmented networks slowed the transmission of infection far more than information (Figs. 2 and 3). Additionally, within these more fragmented, highly modular networks, a higher density of connections led to an increasing gap between the time take for 75% of individuals to become infected and informed (Fig. 3). Qualitatively identical patterns were apparent when considering the difference in spread speeds between modular and random networks (Fig. 4); both information and disease spread more slowly in modular networks than in random networks. Modular networks resulted in a greater reduction of disease transmission speed than information transmission speed when compared to random networks, under most conditions.

**Figure 2:**
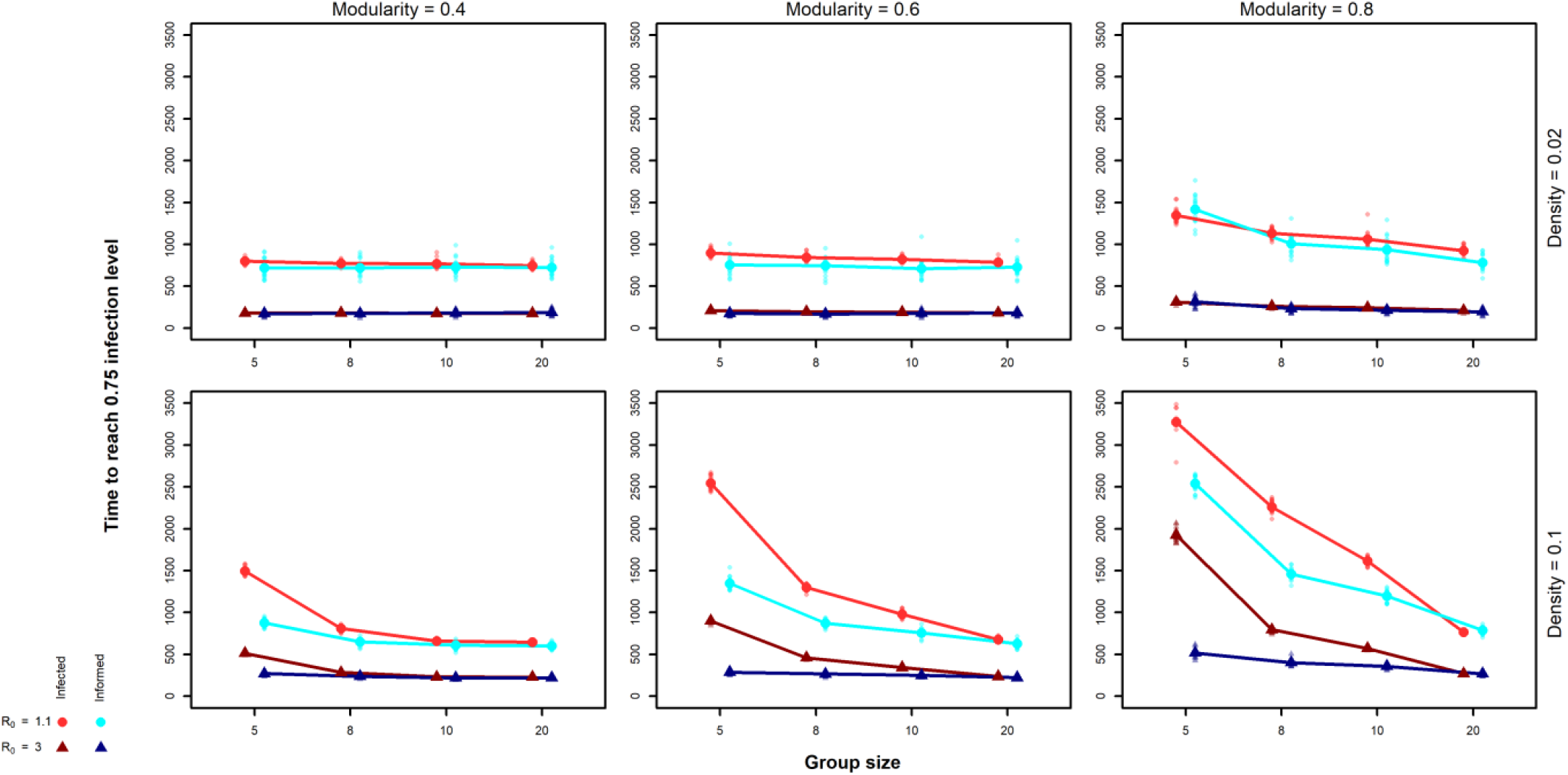
Overview of time taken for disease (red) and information spread (blue) to infect 75% of nodes in a network. Smaller datapoints are raw data while large datapoints are means. Results are shown for different levels of network modularity (column), density (row) and R_0_ (symbol and colour). Note that, for illustrative purposes, we include only a subset of group sizes and other parameter values in this figure, while the full dataset is visualised in supplementary figure 2.

**Figure 3:**
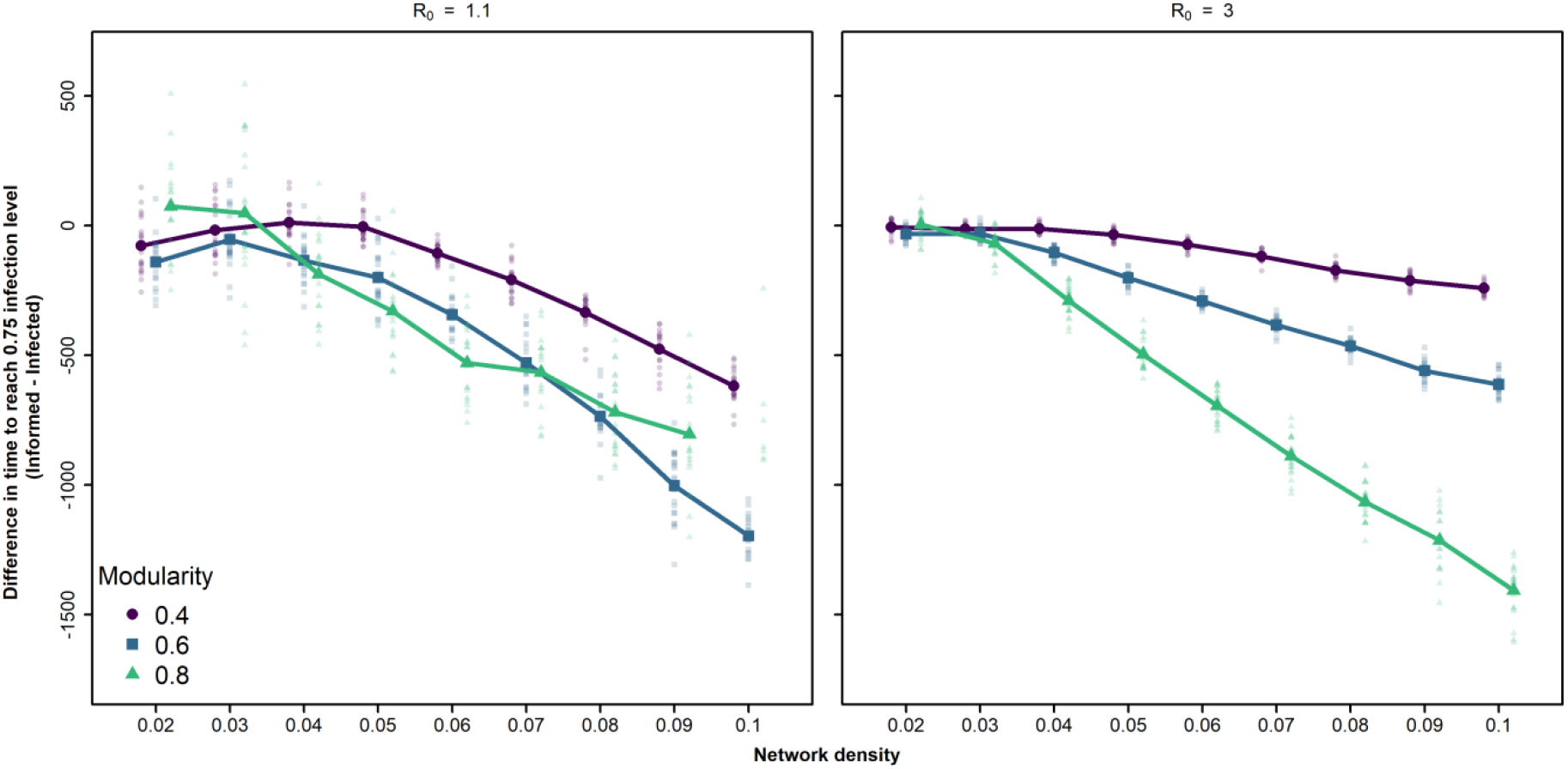
Effect of network density on difference in time taken for disease and information spread to infect 75% of nodes in a network with small groups. The Y-axis shows the time taken for disease to infect 75% of nodes subtracted from the time taken for information to inform 75% of nodes. Positive values on the Y-axis therefore indicate information taking longer to reach this level of infection than disease, while negative values on the Y-axis indicate disease taking longer. Smaller datapoints are raw data while large datapoints are means. Results are shown for different levels of R_0_, network modularity and density, with group size fixed at 5 and population size fixed at 200. Means for parameter combinations where

**Figure 4:**
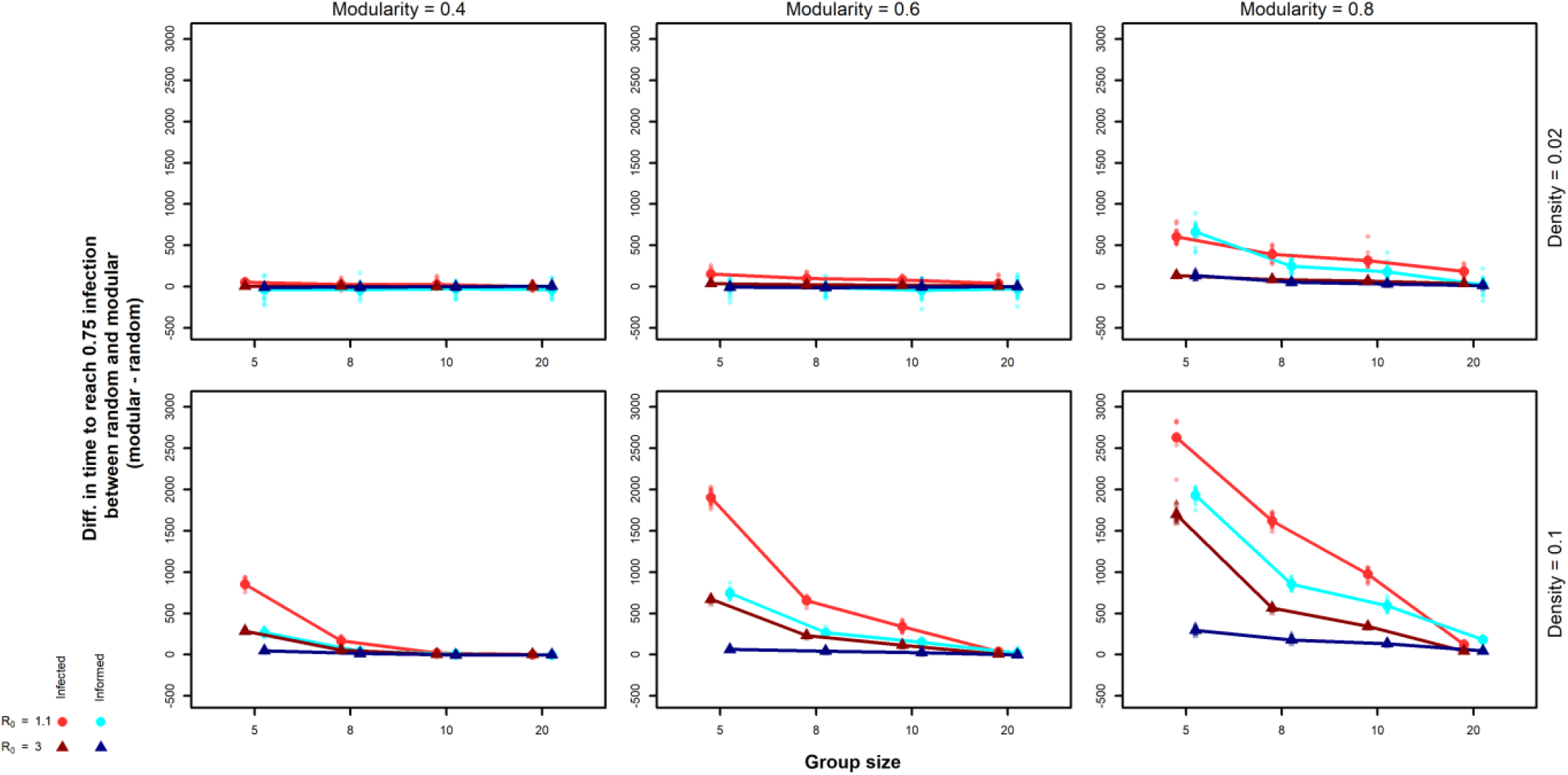
Overview of the difference in time taken for disease (red) and information spread (blue) to infect 75% of nodes in a network between modular and random networks. The Y-axis shows the time taken for a transmission process to reach 75% of nodes in a random network subtracted from the time taken for a transmission process to reach 75% of nodes in a modular network derived from that random network. Positive values therefore indicate slower spreading in modular networks, negative values faster spreading. Smaller datapoints are raw data while large datapoints are means. Results are shown for different levels of network modularity (column), density (row) and R_0_ (symbol and colour). Note that we include only a subset of group sizes and other parameter values in this figure, for a figure showing full dataset see supplementary figure 3.

### The impact of network modularity and fragmentation on transmission rate

In general, it took longer for 75% of the population to become infected and informed in more modular networks (Q_rel_ of 0.6 and 0.8), but only when group sizes were small and more so when transmissibility was lower and network density higher (Fig. 2, Fig. S2). When transmissibility and modularity were high (R_0_=3 and Q_rel_≥0.6) then the time taken for 75% of the population to be informed remained effectively unchanged from that in random networks, while time taken for 75% to be infected increased substantially compared with the random networks (Fig. 4). With lower modularity (Q_rel_ of 0.4), only the smallest group sizes were able to slow transmission, and only when transmissibility was very low. There were 19 dense, highly modular networks with small group sizes where disease entirely failed to spread within 3500 timesteps when R_0_ was low (1.1 or 1.25).

### The Influence of network density and transmissibility on the gap between information and infection spread

The time difference between 75% of the population becoming informed via conformist learning and 75% being infected increased with network density (Fig. 3), although the nature of this relationship depended on both network modularity and transmissibility. When transmissibility was higher, the difference between the rate of information spread and rate of disease spread increased consistently as the modularity of the network increased (Fig 5). The extent of the increase in difference depended on the density of connections in the network. However, when transmissibility was lower the relationship between modularity and difference in transmission speeds was more complicated. First, for networks with intermediate modularities (Q_rel_ of 0.4 or 0.6), the difference in the time taken for 75% of the population to be reached by the two transmission processes showed a (slightly) humped relationship, with the difference in transmission rate being minimised (i.e. close to zero) at low- to intermediate network densities, with disease spreading faster than information in some repeats of the simulation. Second, at high network densities the difference in transmission rates was higher for intermediate modularities (Q_rel_=0.6) than high modularities (Q_rel_=0.8), although it should be noted here that for high modularities disease failed to infect 75% of the population in some runs (see above). Consequently, this result should be seen as a lack of evidence of a difference between intermediate and high modularities rather than a quantitative difference between the two.

**Figure 5:**
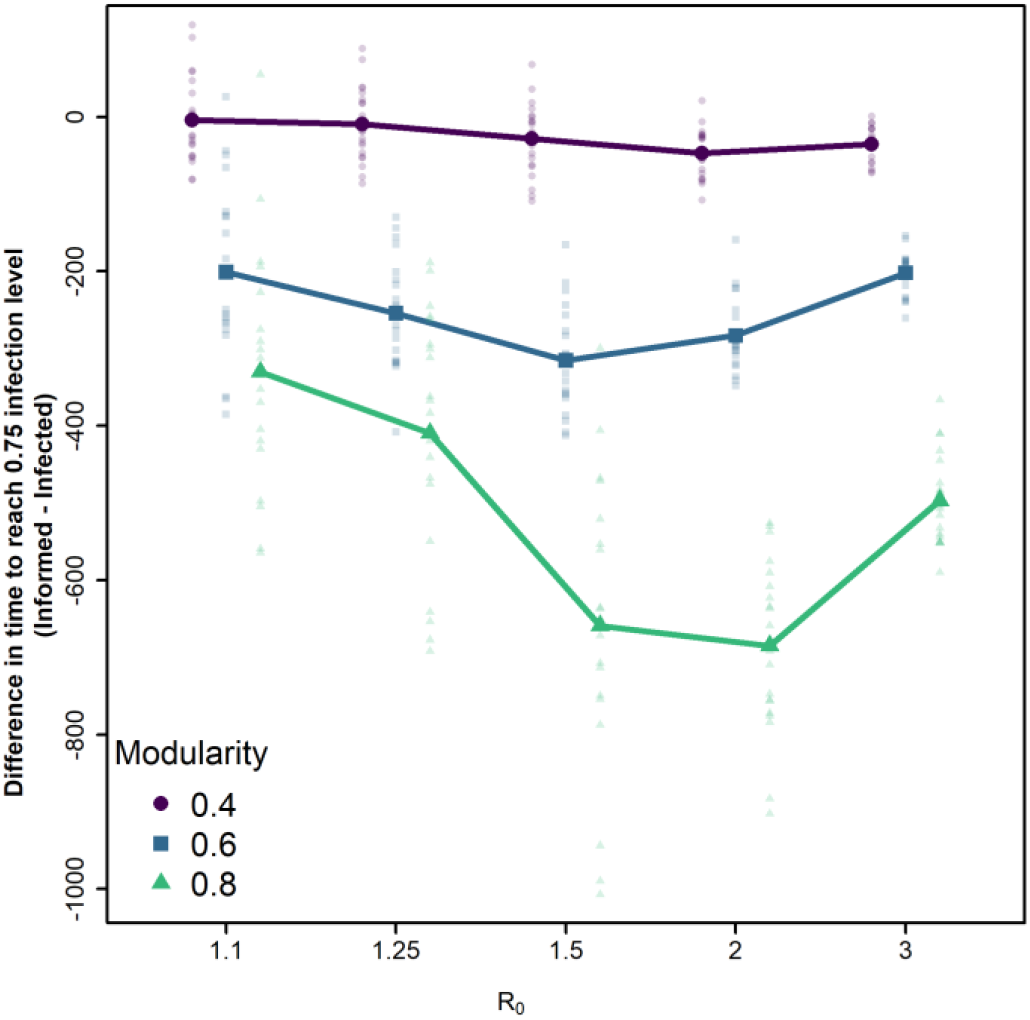
Effect of transmissibility on the difference in time taken for disease and information spread to infect 75% of nodes in a network with small groups. The Y-axis shows the time taken for disease to infect 75% of nodes subtracted from the time taken for information to inform 75% of nodes. Positive values on the Y-axis therefore indicate information taking longer to reach this level of infection than disease, while negative values on the Y-axis indicate disease taking longer. Smaller datapoints are raw data while large datapoints are means. Results are shown for different levels of network modularity, with network density fixed at 0.05, group size fixed at 5 and population size fixed at 200.

For networks with intermediate or high modularity the difference in the rate of transmission was maximised for intermediate values of transmissibility (Fig. 5), resulting in minima at R_0_=1.5 for networks with Q_rel_=0.6 (intermediate modularity) and R_0_=2 for networks with Q_rel_=0.8 (high modularity) among the parameter values we tested.

## Discussion

We show that both modularity and group size are important in mediating the balance between pathogen and information spread when individuals use conformist social learning. In our simulations it took longer for three quarters of individuals to become either informed or infected in modular networks with small group sizes, but not when group sizes were larger. In more fragmented modular networks, the rate of information transmission by conformist learning slowed less than that of infectious disease, resulting in information spreading faster than infection. Because the impact of network fragmentation was smaller, the rate of information transmission differed less between modular and random networks than the rate of disease transmission.

Our simulations clearly demonstrate that modularity can, but does not always, slow the rate of spread of infection, supporting the findings of previous work (Salathé and Jones 2010a; Griffin and Nunn 2012; Nunn et al. 2015; Sah et al. 2017b). Modularity decreased the rate of disease spread most when there was a higher density of network connections and the size of groups was smaller.

The smaller impact of modularity at low network densities is likely because scarce opportunities for transmission in the network as a whole become a more limiting factor than which particular individuals are connected. The role of group size is similar to that found by Sah et al. (2017b), who showed that both modularity and network fragmentation are important in reducing outbreak size in modular networks. Our findings re-emphasise how size of groups is critical in shaping the role of modularity in epidemic dynamics. The impact of modularity on how quickly 75% of our simulated populations became infected was greatest when transmissibility was lower, as would be expected from previous research (Sah et al. 2017b; Rozins et al. 2018). For highly transmissible pathogens and intermediate modularities, even small group sizes were unable to meaningfully slow the spread of disease.

We found similar general patterns for the spread of information via a conformist learning rule. However, while modularity and fragmentation affected information transfer in a similar direction to disease spread, modular networks did not slow down transmission of information as much as the spread of infection compared with random networks. This was especially the case for networks with intermediate modularities and high densities. In general, we did not find support for modularity leading to an increase in the rate of information transmission in our models, contrary to some previous findings (Sah et al. 2017b; Romano et al. 2018). In our results, there was no consistent tendency for information to spread faster in modular networks than in random networks. However, at low network densities and intermediate modularities (Q_rel_ of 0.4 or 0.6), some individual simulation runs found faster information spread in modular than random networks. It is possible, therefore, that modularity might facilitate faster information spread in our modelling framework in parameter space we did not explore in this study, such as a steeper threshold for conformist learning (González-Avella et al. 2011; Nematzadeh et al. 2014). In a previous study that included empirically collected primate networks, network efficiency (a proxy for how efficiently a network exchanges information) was shown to have a relatively weak relationship with modularity that typically peaked at smaller levels of modularity than those investigated here (Romano et al. 2018). Together with our findings, this suggests that the fastest rate of information spread happens in these less modular networks, while more modular networks favour the rate of information transmission relative to the spread of infection.

Conformist social learning has been reported in taxonomically diverse species (e.g. primates:van de Waal et al. 2013; birds: Aplin et al. 2015; insects: Danchin et al. 2018), and so the difference between information and infectious disease transmission shown in our models has ecological and evolutionary significance. Social groups with modular social structure and sufficiently small sub-groups will impede the transmission of infection while having much less impact on how quickly beneficial information spreads. The fact that the difference in information and disease spread is substantial only for small sub-groups may contribute to the evolution of hierarchical or multi-level social structures (Cantor et al. 2015; Grueter et al. 2020). In addition, our work suggests capacity for social structure and social learning rules to co-evolve. In species with social groups that have modular social structures (Nunn et al. 2015) such as fission-fusion societies (Lehmann et al. 2007; Ramos-Fernández and Morales 2014), we would predict conformist social learning to be more favoured than in species without these structures or sub-groupings, as network fragmentation is integral to maintaining higher rates of information spread. Finally, our findings will also be relevant to the spread of information directly related to infectious disease (Funk et al. 2009). When sub-groups and small networks have intermediate modularity, information about the presence of disease will be able to spread faster than the infection itself, helping to enhance behavioural or social immunity (De Roode and Lefèvre 2012; Meunier 2015). The difference will be greatest when the density of connections in a group is higher, as might be expected in a social insect colony for example, or when comparing smaller human communities (higher network fragmentation) to large cities (networks likely still modular but with much bigger groups).

Our results also complement recent modelling and empirical research suggesting modular networks have minimal effect on the spread of social information (Cantor et al. 2021; Laker et al. 2021). For example, Laker et al. (2021) showed that the transmission of novel foraging information was not impacted by the modularity of network structure in domestic fowl chicks *Gallus gallus domesticus*. If we assume conformist learning, our models show that for the modularities investigated in their study (Q=0.63-0.73), we would predict either no increase or a relatively small increase in the rate of learning, depending on how easy information is to acquire. The higher the likelihood of learning at each time-step, the less impact even small sub-groupings have on how quickly information spreads. However, information that spreads according to other social learning rules may respond differently to more modular social structures.

There are a number of limitations to our current model that would benefit from further development. We have only considered binary, unweighted networks at each time-step (a social connection either exists or not). In many situations social connections will vary in strength between different individuals, which could influence the likelihood of transmission, especially of information (Valsecchi et al. 1996; Swaney et al. 2001). Similarly, we have only considered one type of social learning rule, while in reality individuals may use other rules (Kavaliers et al. 2005; Kendal et al. 2015; e.g. biasing learning towards kin or familiar individuals; Boogert et al. 2018) or a combination of multiple rules. A more comprehensive exploration that incorporates heterogeneity in how individuals learn would provide additional insight into how information spread might be influenced by group size and fragmentation. Our model also uses a static network, with no changes over time. It is likely that network dynamics will influence how information and infection spread through a population, with both transmission types potentially causing changes in network structure(Croft et al. 2011; Kulahci et al. 2018; Stockmaier et al. 2020). For example, individuals may be removed from the network due to infection, or the position of individuals within a network might be expected to change when they are infected (Lopes et al. 2016; Stockmaier et al. 2020) or become informed (Kulahci et al. 2018). Future models that incorporate these additional aspects of social structure are likely to provide further insights into the patterns demonstrated here.

Overall, our research supports the idea that modular social structure can help mediate the trade-off between mitigating the spread of infection and enhancing the spread of useful information in animal groups (Evans et al. 2020). However, the extent of these differences in transmission is mediated by the size of groups and network density. We also demonstrate that conformist social learning has the potential to play an important role in helping information spread faster than disease in animal societies, especially in fission-fusion or hierarchical societies characterised by small subgroups. Overall, our results highlight the importance of considering network fragmentation in addition to modularity when studying the ecological consequences of animal social network structure and their implications for the evolution of sociality.

## Acknowledgements

We would like to thank Coby Thompson-Knight for their contribution to preliminary work related to this study. We would also like to thank Barbara König, Jordi Bascompte and Nina Fefferman.

## Declarations

### Data availability

Simulation results are available in the supplementary materials

### Code availability

Simulation code is available in the supplementary materials

### Authors’ contributions

All authors conceived the study. JCE and MJS coded and ran the simulations and wrote the manuscript. All authors discussed the results and contributed to the final manuscript.

