## Supplementary figures and images for "Group size and modularity interact to shape the spread of infection and information through animal societies"

### supplementary figure 1

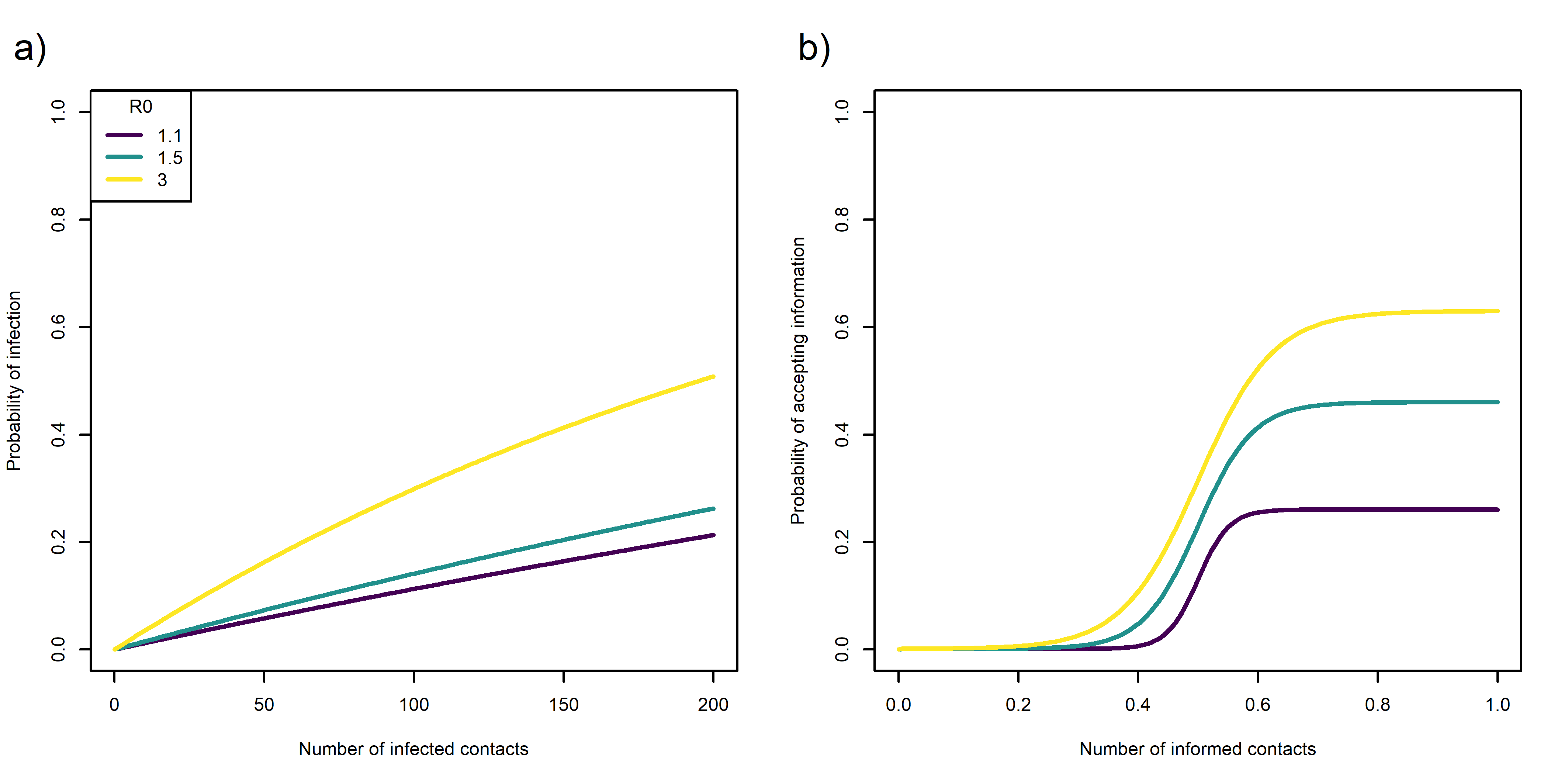

### supplementary figure 2

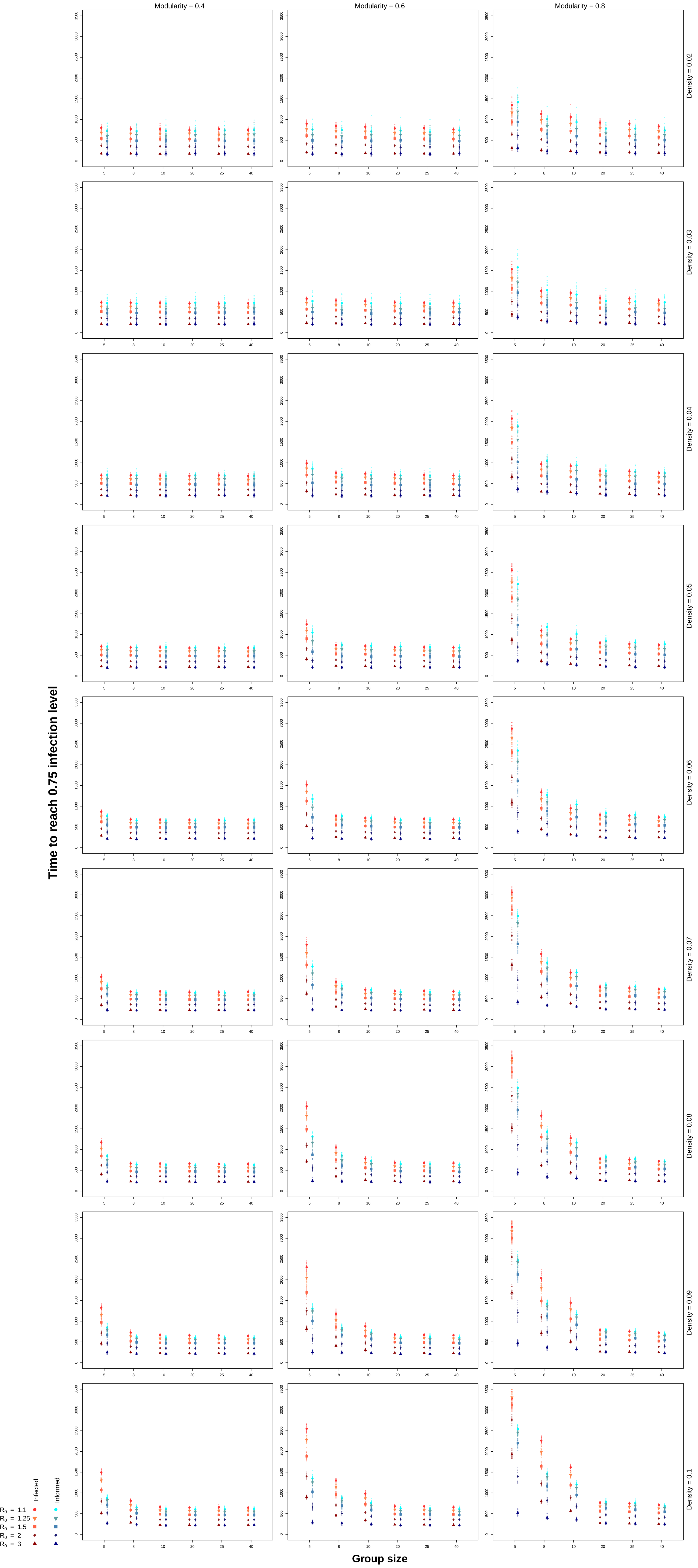

### supplementary figure 3

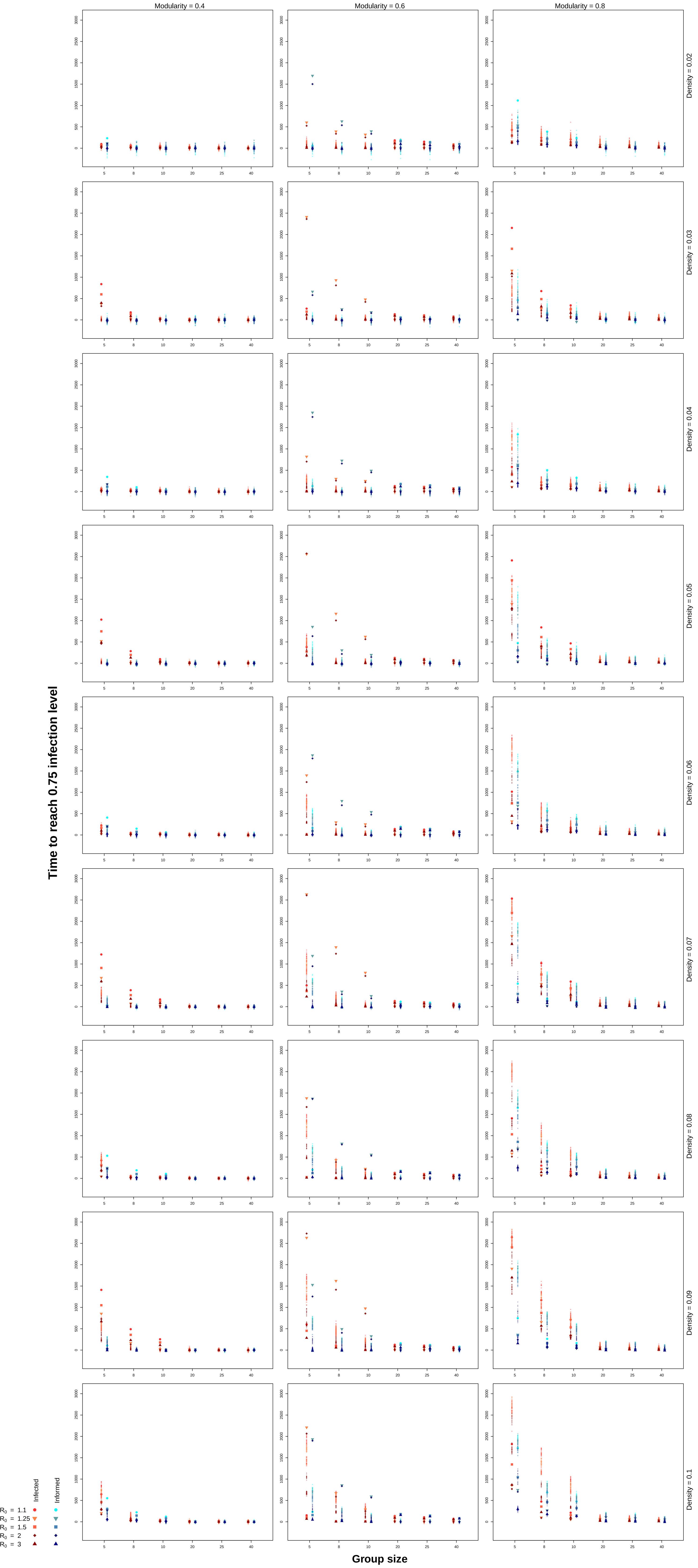
